# Comparative analysis of culture- and ddPCR-based wastewater surveillance for carbapenem-resistant bacteria

**DOI:** 10.1101/2024.06.20.597808

**Authors:** Siyi Zhou, Esther G. Lou, Julia Schedler, Katherine B. Ensor, Loren Hopkins, Lauren B. Stadler

## Abstract

With the widespread use of last-resort antibiotics, carbapenems, clinical reports of infections associated with carbapenem-resistant *Enterobacterales* (CRE) have increased. Clinical surveillance for CRE involves susceptibility testing and/or whole genome sequencing of resistant isolates, which is laborious, resource intensive, and requires expertise. Wastewater surveillance can potentially complement clinical surveillance of CRE, and population-level antibiotic resistance (AR) surveillance more broadly. In this study, we quantitatively and qualitatively compared two widely used methods for AR wastewater surveillance: (1) a culture-based approach for quantifying carbapenem-resistant bacteria and (2) a digital droplet PCR (ddPCR) assay targeting five major carbapenemase genes. We developed a multiplexed ddPCR assay to detect five carbapenemase genes and applied it to wastewater samples from three sites over 12 weeks. In parallel, we quantified carbapenem resistant bacteria and carbapenemase-producing bacteria using culture-based methods. We assessed associations between the concentrations of carbapenemase genes and resistant bacteria. Although both approaches showed similar trends in the overall abundance of dominant carbapenem-resistant bacteria and genes, there were weak correlations between the quantitative levels of resistance. Nanopore sequencing of the resistome of the carbapenem-resistant bacteria revealed that discrepancies arose from differences in the sensitivity and specificity of the methods. This study enhances our understanding of the application of wastewater surveillance in tracking carbapenem resistance and highlights how method choice impacts the results from AR wastewater surveillance.

## 1. Introduction

Antibiotic resistance (AR) has emerged as a major global public health threat ^1^. Infections caused by multidrug-resistant bacteria are on the rise, leading to elevated mortality rates, extended hospital stays, and increased healthcare costs ^2^. Currently, the world is experiencing a silent pandemic of AR ^3^. A recent study revealed that more than 1.2 million people died as a direct result of antibiotic resistant infections in 2019 ^4^. Without intervention, annual deaths could reach 10 million by 2050 ^5^.

Multidrug-resistant *Enterobacterales* have become a significant public health concern due to the inappropriate use of antibiotics coupled with the widespread prevalence of *Enterobacterales* in both healthcare and community-associated infections ^6^. Carbapenems are considered last-resort medication for serious multidrug-resistant infections ^7^. However, the prevalence of carbapenem-resistant *Enterobacterales* (CRE) is rising globally ^8–10^. CRE was named one of three critical priority pathogens for new antibiotics development by WHO in 2017 ^11^ and remain one of the top five most urgent public health threats in 2019 by the U.S. Centers of Disease Control and Prevention ^12^. Carbapenem resistance in *Enterobacterales* may be conferred due to the production of carbapenemase enzymes, which are encoded by a diverse group of genotypes and often encoded on plasmids ^13^. Globally, the most common types of carbapenemases include *Klebsiella pneumoniae carbapenemases* (KPC), oxacillinase (OXA)-48-like β-lactamases, and metallo-β-lactamases such as New Delhi metallo-β-lactamases (NDM), active-in-imipenem family of carbapenemases (IMP), and Verona integron-encoded metallo-β-lactamases (VIM) ^14^. Carbapenem resistance in CRE may also arise from the production of extended-spectrum β-lactamases (ESBLs) and/or AmpC enzymes, combined with changes in permeability due to the loss of outer membrane porins or the overexpression of efflux pumps ^15,16^. Additionally, some *Enterobacterales* have intrinsic resistance to imipenem ^17,18^.

Local epidemiology of AR can guide treatment strategies and alert healthcare providers to new forms of resistance ^10^. Traditional clinical surveillance, including susceptibility testing and whole genome sequencing (WGS) of resistant isolates, is crucial for optimizing patient care and controlling the spread of resistant pathogens. For instance, a study investigating patient samples from South Texas between 2011 and 2019 found that infections caused by non-carbapenemase-producing *Enterobacterales* were associated with increased prior antimicrobial use and more emergency visits ^8^. WGS of resistant isolates has proven effective in linking phenotypic and genotypic profiles ^19^. For example, genomic surveillance was used to analyze the specific carbapenem-resistance determinants of clinical isolates of *Pseudomonas aeruginosa* in Pakistan. The findings revealed that the acquisition of multiple resistance mechanisms correlated with increased levels of imipenem resistance, emphasizing the combinatorial resistance mechanisms of carbapenemase-production and *oprD* mutations ^20^. However, these methods are costly, labor-intensive, and inefficient for large-scale population monitoring. Also, given that carbapenem-resistant bacteria are not confined to healthcare environments and may proliferate in community settings, traditional clinical surveillance may be biased toward patient samples collected in healthcare facilities.

Wastewater surveillance has the potential to detect the spread of carbapenem resistance genes and bacteria throughout communities. Previous studies have highlighted the potential of wastewater monitoring for tracking AR at the population level ^10,19,21^. Two common methods used for wastewater surveillance of carbapenem resistance are: (1) quantifying specific carbapenemase-encoding genes via quantitative PCR (qPCR) or digital droplet PCR (ddPCR) ^22,23^; and (2) culturing carbapenem-resistant colonies in wastewater ^24–26^. Culture-based methods focus on viable bacteria enriched with selective media, and when paired with molecular tests like conventional PCR or WGS, yield insights into phenotype-genotype relationships ^19^. The selective culturing of resistant isolates can reveal rare, low-abundance resistant organisms, and sequencing them can provide potential mechanisms of resistance ^27^. Culture-based methods, however, likely miss the majority of bacteria in sewage, given that a high proportion of environmental microbes cannot be cultured under conventional laboratory conditions ^28–30^. Molecular methods such as qPCR or ddPCR can overcome some limitations of culture-based methods. These techniques enable broader detection and quantification of target genes present in culturable organisms, extracellular DNA, and viable but non-culturable organisms in wastewater. However, PCR-based techniques have several limitations. Firstly, such methods require prior knowledge of target genes before conducting experiments. Moreover, these methods cannot differentiate between DNA from live and dead cells, which may lead to an overestimation of pathogen loads or microbial abundance in samples. Although PCR can identify genetic material, it is unable to determine whether genes are actively expressed or have an effect on an organism’s phenotype. Additionally, PCR-based methods are susceptible to contamination and require rigorous experimental controls and cleanliness to ensure accurate results. Lastly, by solely tracking ARGs, it is challenging to estimate antibiotic resistant bacteria (ARB) abundance in wastewater, as the copy numbers of ARGs can vary in different bacteria ^31^.

Current wastewater surveillance of AR is hampered by a lack of understanding how method choice impacts the data and trends observed, and thus the ultimate actionability and interpretation of the information generated. Quantitative measurements of AR targets could be used to establish excessive pathogen and/or gene abundances, as well as determining the appropriate thresholds for triggering alerts in response to wastewater detections, as has been done for viral wastewater surveillance ^32–34^. However, quantification methods for AR in wastewater, and specifically the comparability between culture-based and molecular-based detection methods have not been established. Comparative quantitative studies are scarce, with only a handful of investigations attempting to address this critical need. Flach et al. found that certain carbapenemase genes including *bla*_OXA-48-like_, *bla_NDM_*, and *bla_KPC_* were associated with the abundance of carbapenemase-producing *Enterobacterales* in hospital sewage, but not other carbapenemase genes ^10^. Rocha et al. also compared qPCR and culture-based methods for quantifying antibiotic resistance in wastewater but did not include carbapenem resistance ^35^. Further research is needed to assess associations between quantitative measures of antibiotic resistance, and particularly priority pathogens such as CRE, in community wastewater. This lack of comparative data complicates cross-study comparisons of AR wastewater surveillance data. In this study, we directly compared two approaches: 1) a culture-based method for carbapenem-resistant bacteria quantification ^36^, and 2) a ddPCR assay for five representative ARGs encoding for carbapenemase-production (CP). We then examined associations between the quantitative results yielded from each approach generated from surveillance of samples collected from three wastewater treatment plants over 12 weeks. To further investigate the potential mechanisms responsible for the carbapenem-resistant phenotypes, we used Nanopore sequencing to characterize the resistance profile of the carbapenem-resistant bacteria isolated using the culture-based method.

## 2. Methods

### 2.1 Wastewater sample collection, concentration, and DNA extraction

Weekly wastewater samples were collected from three wastewater treatment plants (WWTPs, 69^th^, UB and WD, respectively) in Houston, Texas, between August 22 and November 7, 2022. Untreated wastewater was collected using a refrigerated autosampler that collected a time-weighted composite sample of raw wastewater by sampling every hour over a 24-hour period from the influent of the WWTPs. Wastewater samples, each with a volume of 50 mL, underwent centrifugation at 4,100 g for 20 minutes at 4°C. The resulting pellets were processed immediately for DNA extraction and stored at −20°C for analysis. DNA extraction was performed using the Chemagic™ Prime Viral DNA/RNA 300 Kit H96 (Chemagic, CMG-1433, PerkinElmer). Detailed concentration and DNA extraction procedures are provided in the Supplementary materials (Materials and Methods) following the Environmental Microbiology Minimum Information (EMMI) Guidelines ^37^.

### 2.2 Culture-based methods

For the quantification of carbapenem resistant (CR) bacteria in wastewater, a stepwise procedure was applied, utilizing membrane filtration and selective media supplemented with a carbapenem ^36^. Briefly, presumptive CR bacteria were screened using membrane filtration on selective media with an intermediate concentration of ertapenem sodium (Sigma®). Subsequently, individual colonies from the presumptive test were cultured on media containing a higher ertapenem concentration to eliminate false positives and intermediate-resistant colonies. Carbapenemase production of each confirmed CR colony was finally assessed using the Blue-Carba test.

Specifically, membrane fecal coliform Agar (mFC, Difco® with 0.01% (w/v) Rosolic Acid (Difco®)) was prepared according to manufacturer instructions. For the quantification of presumptive CR bacteria, mFC Agar was supplemented with ertapenem sodium at a final concentration of 1 μg/mL. This concentration served as an intermediate susceptibility level, aligned with the standards provided by the Clinical and Laboratory Standard Institute (CLSI) ^38^. To ensure the maximum activity of the carbapenems, mFC selective media was prepared and dispensed into 60 mm diameter Petri plates the day before use, then stored at 4 °C in the dark. Wastewater samples were serially diluted, and 1 mL of the appropriate dilution was filtered through a 0.45 μm nitrocellulose sterile membrane filter. The membrane filters were placed on mFC Agar plates with and without ertapenem. All membrane filtration procedures were performed within six hours of wastewater sample collection. Cultures were incubated for 24 hours at 37 °C after which colonies were enumerated to calculate the number of presumptive CR bacteria. Subsequently, the presumptive CR colonies were counted and transferred onto Mueller Hinton Agar (MHA) media containing an elevated concentration of 2 μg/mL ertapenem, based on the CLSI breakpoint. The cultures were then incubated at 35.0 °C for 24 hours. Growth on the selective MHA media was recorded as positive confirmation and these colonies were enumerated to calculate the number of confirmed CR bacteria. Next, the confirmed CR colonies were subjected to further evaluation for carbapenemase production using the Blue Carba test ^39^. In the Blue Carba test, CR colonies were introduced into an imipenem (Sigma®)-supplemented solution containing bromothymol blue as a pH indicator and then incubated for a duration of two hours at 35°C. A color change from blue to yellow/green signified a positive result. Control experiments were conducted using *K. pneumoniae* BAA-1705 (carbapenem-resistant) and wild-type *E. coli* MG1655, confirmed to be susceptible to carbapenem. All experiments were performed in triplicate.

### 2.3 Multiplexed ddPCR assay for the quantification of carbapenemase genes

A multiplex ddPCR assay was designed to detect five common carbapenemase groups: *bla*_NDM_, *bla*_KPC_, *bla*_VIM_, *bla*_IMP_ and *bla*_OXA-48-like_. The primers and probes were designed using RefSeq sequences for the five carbapenemase-encoding genes and their variants, sourced from the National Center for Biotechnology Information (NCBI) as of June 2022. A total of 123 RefSeq sequences were retrieved for the KPC carbapenem-hydrolyzing class A beta-lactamase gene family, 43 for the NDM subclass B1 metallo-beta-lactamase gene family, 78 for the VIM subclass B1 metallo-beta-lactamase gene family, 48 for the blaOXA-48-like class D beta-lactamase gene family, and 98 for the IMP subclass B1 metallo-beta-lactamase gene family (Table S3). After applying multiple sequence alignment for each gene family, a conservative sequence was identified by manual inspection primers, and probes were designed for the sequence with Primer3Plus ^40^. The theoretical specificity of primer candidates was checked *in silico* using Primer-Blast ^41^ and selected primers were chosen based on targeted gene coverage, specificity, and thermodynamic properties. To multiplex primer and probe sets, multiple primer analyzer (Thermos Fisher Scientific) was used to check the self- and cross-dimers and calculate melting temperatures for multiplex primers/probes. We further confirmed the specificity of the primer sets by performing PCR on real wastewater samples. Specifically, PCR amplicon product sizes were verified visually using gel electrophoresis and then extracted for Sanger sequencing. In addition, a temperature gradient from 56.2 °C to 62 °C was applied to identify the optimal annealing temperature for the PCR reaction. Details of the multiplexed ddPCR assay are available in the Supplementary materials (Fig. S2). All primers and probes were synthesized by Integrated DNA Technologies (Coralville, IA). The primers and the corresponding probes are listed in the Supplemental materials (Table S4).

### 2.4 ddPCR quantification of carbapenemase-encoding genes

Five carbapenemase-encoding genes, *bla*_NDM_, *bla*_KPC_, *bla*_VIM_, *bla*_IMP_ and *bla*_OXA-48-like_, were quantified via ddPCR. ddPCR was performed using a QX600 AutoDG Droplet Digital PCR System (Bio-Rad) and a C1000 Thermal Cycler (Bio-Rad) in 96-well optical plates. The ddPCR reaction mixture was composed of 5 µL ddPCR Multiplex Supermix (Bio-Rad, Hercules, CA, USA), 0.9 µM of each primer, 0.25 µM of each probe, 10 µL template, and Ultrapure DNase/RNase free distilled water (Thermo Fisher Scientific, Waltham, MA, USA) to reach a final volume of 22 µL. An automatic droplet generator was used to perform the droplet generation (Bio-Rad, Hercules, CA, USA). The droplet-partitioned samples were loaded in a 96-well plate and heat-sealed with a PCR plate heat-seal foil (Bio-Rad, Hercules, CA, USA). The plate was transferred in a thermal cycler (Bio-Rad, Hercules, CA, USA) for PCR amplification, following the cycling procedure in the Supplementary materials (Table S5, S6). The amplified samples were loaded onto a droplet reader (Bio-Rad, Hercules, CA, USA) for data quantification. Five gene fragment standards from Twist Bioscience (South San Francisco, CA, USA) were used as positive controls (Table S7). Two blank samples were included in the concentration and extraction steps and two no-template controls (NTC) were included during ddPCR quantification to check for contamination and serve as negative controls. Detailed method information, including concentration factor calculations, limit of detection (LOD), was described previously and summarized in the Supplementary materials ^42^. ddPCR data was processed using QuantaSoft™ Software version v1.7.4 (Bio-Rad, Hercules, CA, USA).

### 2.5 Nanopore library preparation and sequencing

Presumptive carbapenem-resistant colonies were washed off plates using phosphate-buffered saline (PBS, 1X, Thermo Fisher Scientific) from nine selective agar plates that contained samples from three WWTP sites. Genomic DNA was extracted from the mixture of colonies from each plate using a Quick-DNA HMW MagBead Kit (Zymo Research, USA) following the manufacturer’s instructions. The DNA libraries from the nine samples were grouped into three batches. Each batch was then prepared as a multiplexed library using the Native Barcoding Kit 24 V14 (SQK-NBD114.24) following the manufacturer’s instructions. To assess DNA fragment size, we conducted gel electrophoresis. DNA segments exceeding 1 kb in length were manually excised from the agarose gel and retrieved using a Monarch® DNA Gel Extraction Kit (NEB Inc., USA). A Qubit 2.0 Fluorometer and the Qubit (ds)DNA HS Assay Kit (Thermo Fisher Scientific) was used to determine DNA concentrations. DNA quality was checked by a microplate reader (BioTek Take3, Agilent, CA, USA) and only DNA samples that met the purity criteria (OD 260/280 around 1.8 and OD 260/230 between 2.0–2.2) were processed for sequencing. Each of these multiplexed libraries was then sequenced individually on FLO-MIN114 R10.4.1 MinION flow cells over a 12-hour period.

### 2.6 Sequence analysis

Raw reads generated by MinKNOW were base-called and demultiplexed using guppy v6.4.6 (https://community.nanoporetech.com) to yield fastq files. To search for the ARGs, we aligned the base-called reads against the nucleotide sequences of the Comprehensive Antibiotic Resistance Database (CARD, version 3.2.7) using blastn (version 2.14.0+) ^43,44^. We then refined the alignment results by applying strict criteria on alignment length and similarity. Only those alignments that covered over 95% of the ARG length and had a similarity greater than 80% were considered for further analysis ^45^. In cases where regions containing ARGs exhibited more than 80% overlap, only the most accurate ARG match was retained. PlasFlow ^46^(v1.1) was then used to classify the accepted reads into chromosomal and plasmid-associated groups. Adhering to the recommendation of the PlasFlow developer, we filtered out any reads shorter than 1000 bp via Filtlong (v0.2.1, https://github.com/rrwick/Filtlong) since PlasFlow does not perform optimally with short sequences. For taxonomic classification, we used Centrifuge ^47^ (v1.0.4) with the NCBI non-redundant nucleotide sequence database. The categorization outcomes were then visualized using Pavian ^48^. Data analyses and figures were conducted using R.

### 2.7 Statistical analysis

Pearson correlation analysis was conducted to examine the relationship between the colony-forming units (CFUs/mL) across three sites, as well as between colonies (CFUs/mL) and the gene absolute abundance values (gene copy number/mL of the sample) via R. This analysis was based on 36 samples from three WWTPs (69^th^ wastewater influent (n = 12), UB wastewater influent (n = 12), and WD wastewater influent (n = 12)). The analysis focused on the five carbapenemase-encoding genes (*bla*_NDM_, *bla*_KPC_, *bla*_VIM_, *bla*_IMP_ and *bla*_OXA-48-like_), *rpoB*, the sum of the five carbapenemase-encoding genes and the cultured colonies, including total coliforms, presumptive CR bacteria, confirmed CR bacteria and CP bacteria. Since temporal dependence may impact the statistical significance of Pearson correlation, each pair of series was checked for autocorrelation using the Durbin-Watson test in the R package car (Table S8, 9) ^49^.

## 3. Results and Discussion

### 3.1 Both culture- and ddPCR-based methods revealed similar trends in the overall abundances of dominant carbapenem resistance genes in wastewater

We collected a total of 36 influent wastewater samples from three WWTPs over twelve weeks (Fig. 1). Total coliform concentrations (measured in CFU per ml of wastewater) exhibited significant fluctuations, ranging from 5.15 to 6.28 log CFU ml^-1^. Among these, a small portion were categorized as presumptive and confirmed CR bacteria, accounting for 3.33 ± 1.14 % and 2.84 ± 0.89 % of the total coliform counts, respectively. CP bacteria constituted an even smaller fraction, making up 0.44 ± 0.18 % of the total coliform population. Strong correlations between presumptive CR bacteria and confirmed CR bacteria were observed at all three sites (Pearson’s r > 0.95, p < 0.001, Table S8, Fig. S3).

**Figure 1.**
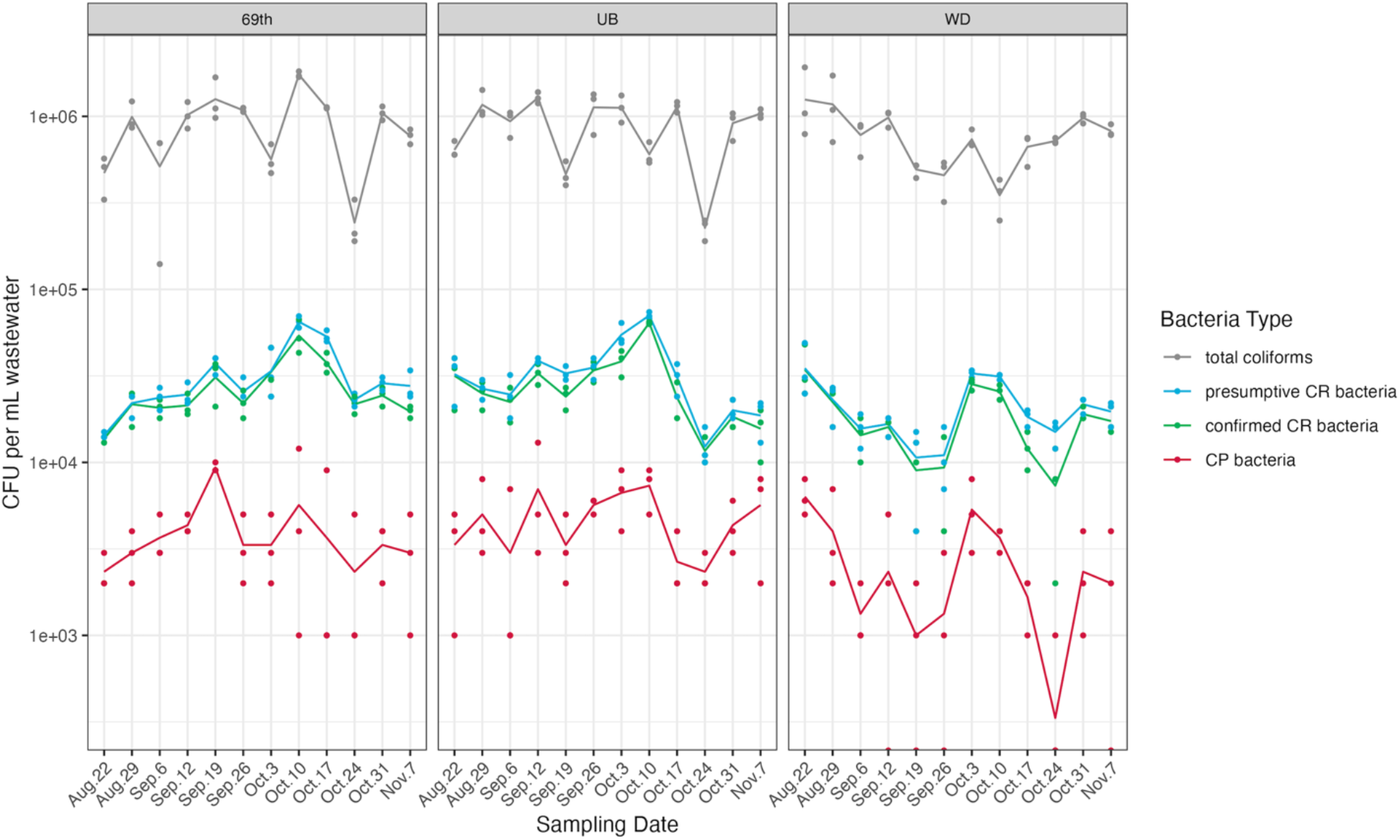
Colony Forming Units (CFUs) of total coliforms, presumptive carbapenem-resistant (CR) bacteria, confirmed carbapenem-resistant (CR) bacteria, and carbapenemase-producing (CP) bacteria across three WWTPs. Each dot represents a replicate, and the lines connect the averages of the triplicates.

In addition to culture-based measurements of carbapenem resistance, we quantified five carbapenemase-encoding genes in the same wastewater samples (Fig. 2). Notably, the patterns of gene absolute abundance in wastewater paralleled that of gene relative abundance (Fig. S4, normalized by *rpoB* genes). Furthermore, the concentrations of these genes varied by site throughout the study, as depicted in Figure S5. Such site-specific distribution patterns for each gene could stem from the distinct service area and geographical positions of the three WWTPs (Fig. S1, Table S1).

**Figure 2.**
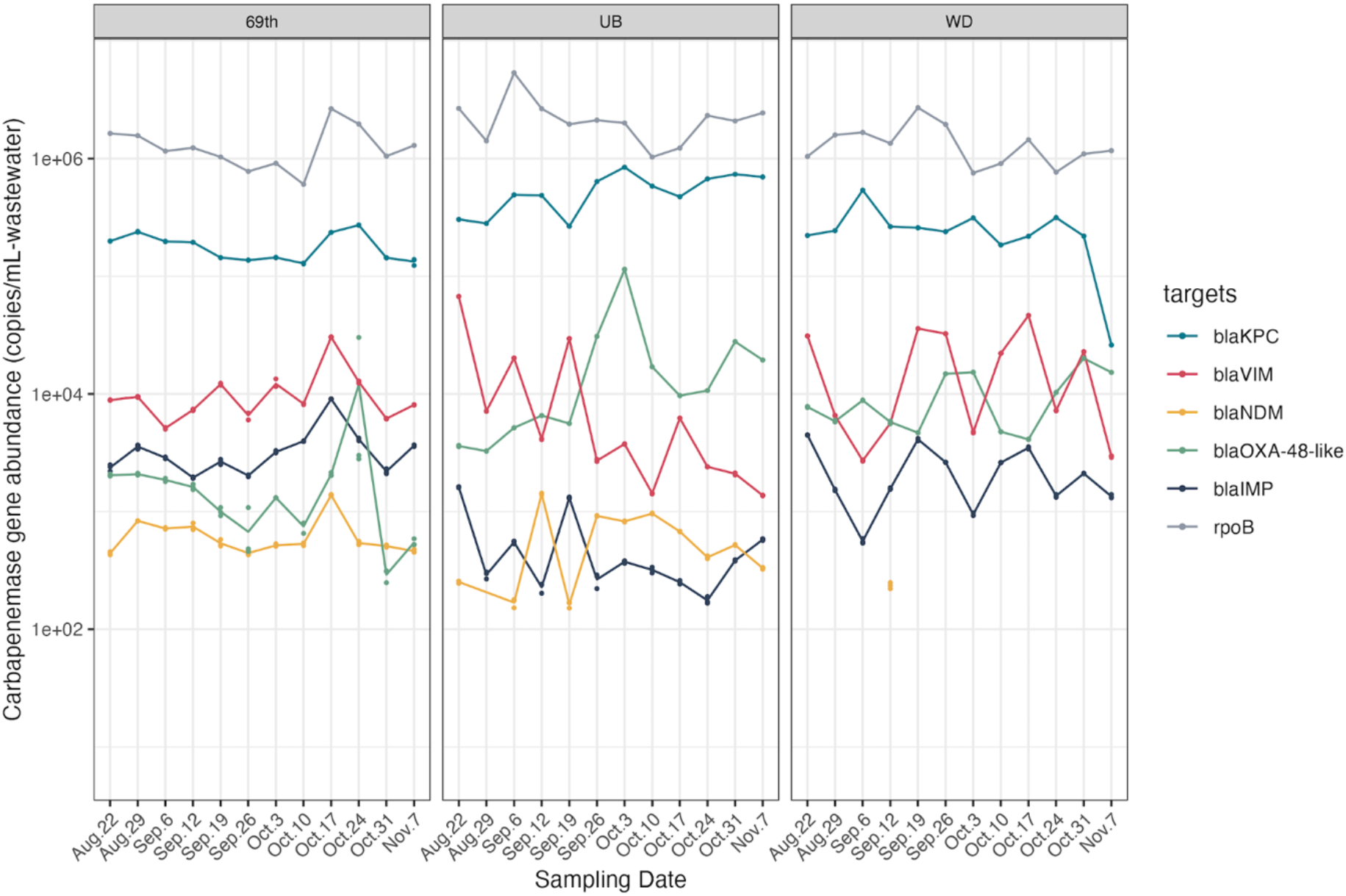
Carbapenemase-encoding gene abundances (copies/mL of wastewater) in the wastewater influent from three WWTPs (69^th^: 69^th^ Street; UB: Upper Brays; and WD: West District). Three replicate samples were analyzed on each sampling date, depicted as dots, and the line connects the mean value of the replicates. Targets below the limit of detection are not shown.

Among these genes, *bla*_KPC_ genes were the most frequently detected and most abundant in all samples, with an average concentration of log₁₀ (5.43 ± 0.29) copies ml ^-1^ of wastewater. This finding was quantitatively consistent with the presumptive CR bacteria which contained high read counts of *bla*_KPC_ family genes as compared with other carbapenemase-encoding genes, revealed by sequencing results. Previous studies reported similar prevalence of *bla*_KPC_ in US wastewater samples, with *bla*_KPC_-containing bacteria making up significant portions of all identified carbapenemase-producing bacteria ^50,51^. Our findings underscore that *bla*_KPC_-containing bacteria are abundant and in US wastewater and ubiquitous in the population.

In contrast, *bla*_NDM_ was present at low concentrations across all sites and frequently was below the detection limit at the WD site. This was also consistent with the sequencing data, as we did not observe any reads for *bla*_NDM_ in the presumptive carbapenem-resistant coliform metagenomes. Moreover, two other carbapenemase-encoding genes (*bla*_VIM_, *bla*_IMP_) were detected at relatively low concentrations via ddPCR, typically one to two orders of magnitude lower than *bla*_KPC_ concentrations. Similar to *bla*_NDM_, neither of these two genes were detected in presumptive CR-coliform metagenomes. PCR-based quantification is more sensitive than culture-based methods for numerous reasons. Culture-based methods favor specific bacterial species or strains that thrive on the selective media under particular conditions. For example, a previous study has shown that *bla*_VIM_ and *bla*_IMP_ genes are often carried in the form of integron gene cassettes and are found commonly in several non-*Enterobacterales* species ^52^. Our ddPCR results of these genes are likely due to the presence of *bla*_VIM_/*bla*_IM_-harboring strains of non-*Enterobacterales* species in the sewage. Further, since culture-based methods are low-throughput, certain non-*Enterobacteriales* species that carry these genes may not have been captured by the culture-based method.

### 3.2 Weak correlations between the quantitative levels obtained using the two different methods were observed

The *rpoB* gene served as a measurement of total bacteria, however, there was not a statistically significant relationship between *rpoB* and total coliform concentrations (Fig. 3, Table S9, r = 0.13, p > 0.999). Moderate correlations were found between both presumptive and confirmed CR bacteria with CP bacteria (r = 0.69, p < 0.001; r =0.72, p < 0.001). Durbin-Watson test for autocorrelation did detect positive autocorrelations between both presumptive and confirmed CR bacetria with CP bacteria, indicating that the statistical significance of the correlations were likely driven by autocorrelation. Additionally, moderate correlations were observed between total coliforms and CP bacteria (r = 0.53, p = 0.047). However, no significant associations between carbapenemase-encoding genes and bacteria counts were observed.

**Figure 3.**
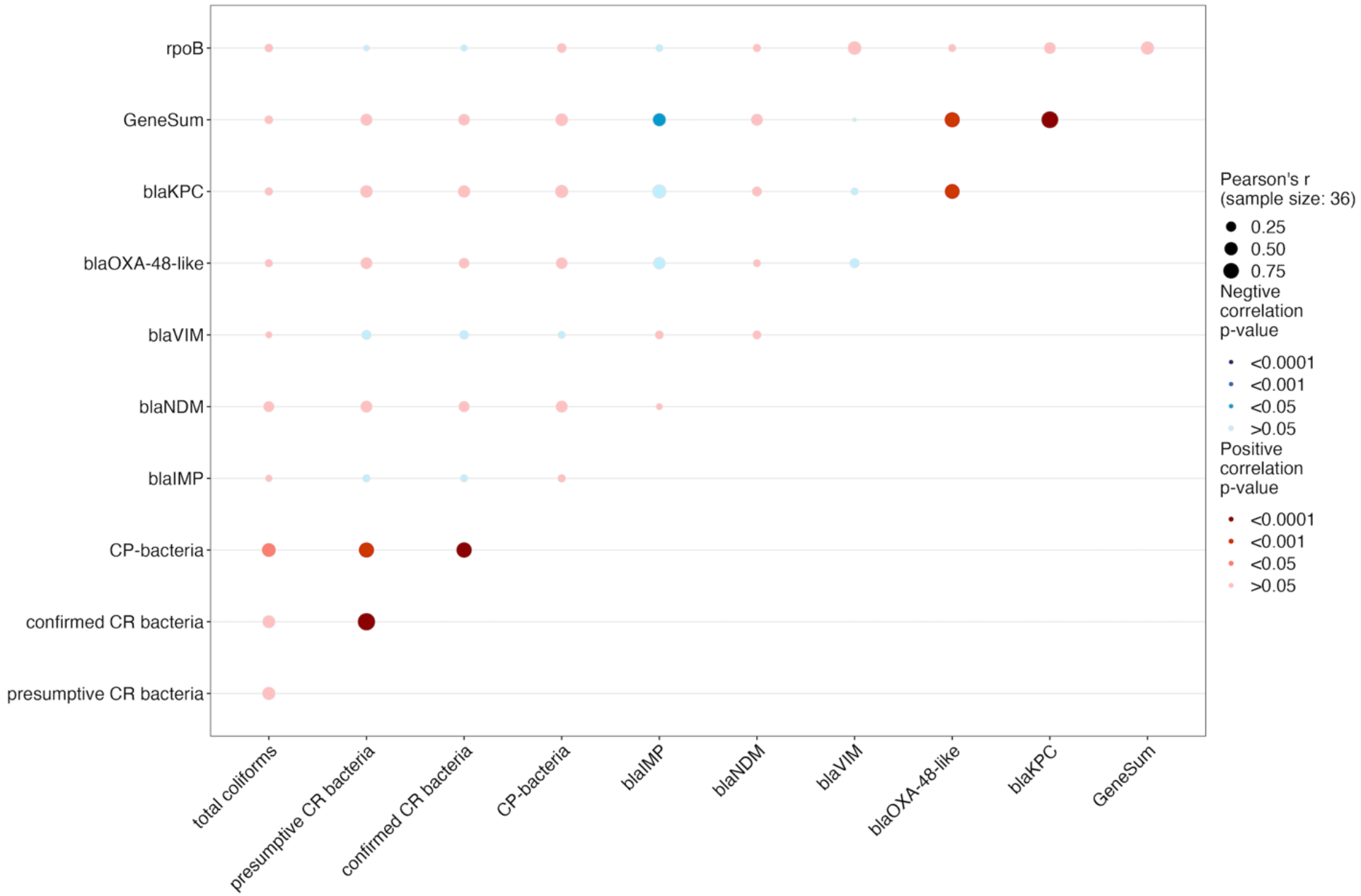
Pearson correlation analysis of concentrations of carbapenemase-encoding genes (*bla*_NDM_, *bla*_KPC_, *bla*_VIM_, *bla*_IMP_ and *bla*_OXA-48-like_) and the cumulative concentration of all five genes (GeneSum), *rpoB* genes, total coliforms, presumptive CR bacteria, confirmed CR bacteria, and CP bacteria. The bubbles represent the correlation between carbapenamase-encoding genes (copy number per ml wastewater) with the concentration of bacteria (CFU per ml wastewater). Bubble size reflects the Pearson correlation coefficient (r), with larger bubbles representing stronger correlations. The color of the bubble indicates the significance level (p): blue signifies a negative correlation, red signifies a positive correlation, and darker shades imply stronger correlation evidence.

There are several reasons why these two methods yielded inconsistent information. First, culture-based methods generally have a significant bias as they exclude non-culturable populations, including many environmental bacteria found in wastewater samples that do not grow on standard culture media, as pointed out by previous studies ^35^. While for ddPCR-based methods, the detection of the target genes is largely agnostic to their host organism and will include genes present in both viable and non-viable bacteria. In our study, the use of mFC agar for the selection of culturable coliforms is widely used due to its cost-effectiveness and simplicity for assessing fecal contamination. However, previous research indicated that carbapenem-resistant bacteria originating from humans in wastewater encompass a broader spectrum of enterobacteria, not just coliforms ^53^. Furthermore, it is important to note that conventional culture-based methodologies tend to favor the proliferation of fast-growing bacterial species, primarily due to the absence of spatial structure in standard laboratory cultivation practices ^54^.

### 3.3 Nanopore sequencing revealed CR mechanisms in cultured samples missed by ddPCR

To understand the potential resistance mechanisms that may have led to the observed carbapenem-resistant phenotypes, and to better understand the weak correlations between the culture-based and ddPCR-based methods, we collected presumptive CR bacteria from three samples from each WWTP site and performed long-read metagenomic sequencing. Nanopore sequencing of presumptive CR bacteria yielded 2.38 Gbps of data with N50 values ranging from 5,880 to 9,580 bp across nine samples from three WWTP sites. In total, 2,956 reads carrying ARGs were identified, with 1,364 of them associated with multidrug resistance. Additionally, we classified the sequencing reads based on their association with either chromosomes or plasmids. Among the detected ARGs, 13 distinct types were identified. Except for multidrug ARGs, plasmids were the dominant carriers, accounting for 767 reads, compared to 557 reads associated with chromosomes (Fig. 4). Beta-lactam resistance genes were the most prevalent, constituting 15.3% of the total ARGs (453 reads). Additionally, ARGs associated with peptides (258 reads, 8.7%), aminoglycosides (194 reads, 6.6%), and fluoroquinolones (189 reads, 6.4%) were abundant, suggesting the potential co-selection of beta-lactamase genes and these types of ARGs using the selective culture media.

**Figure 4.**
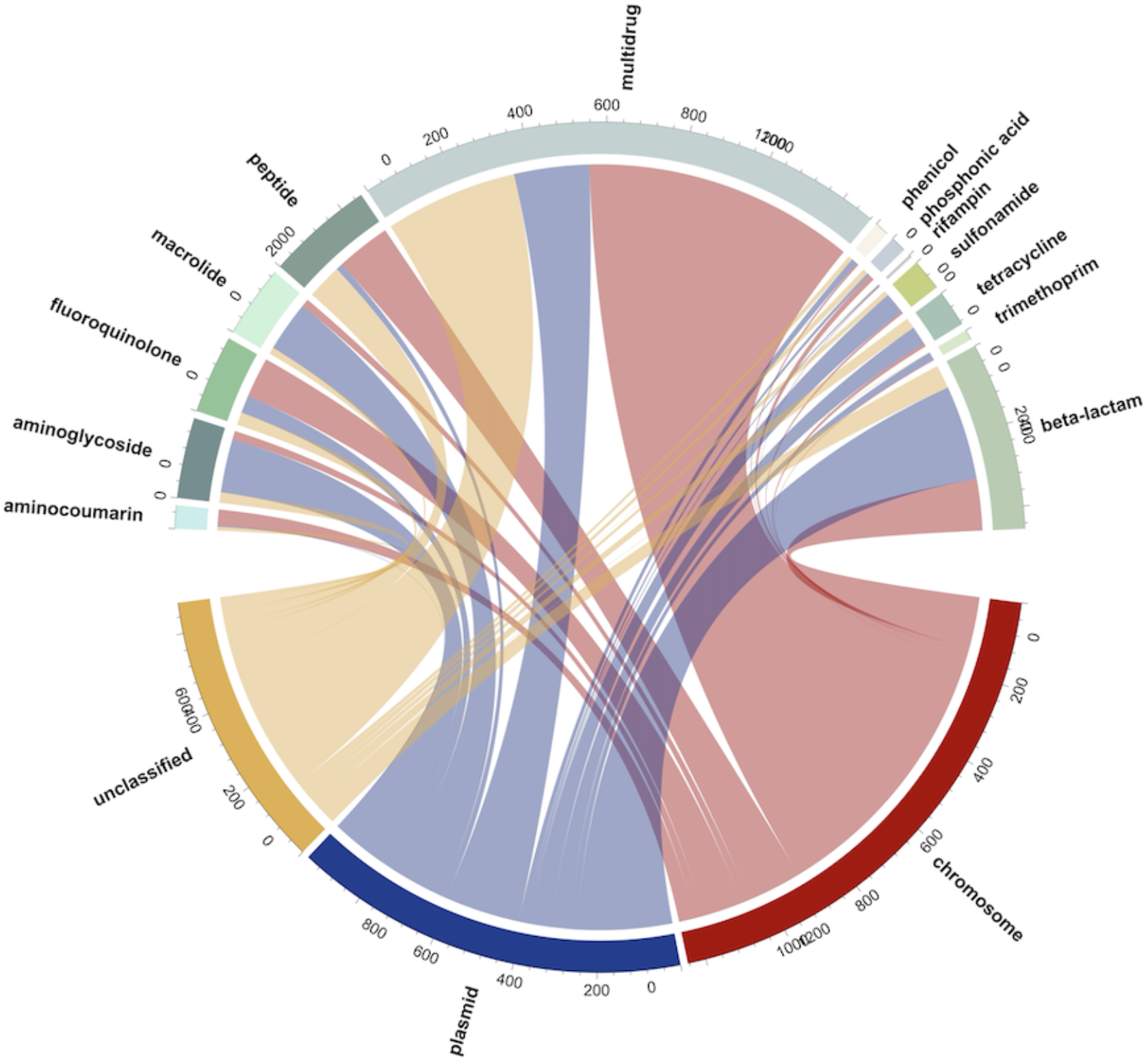
Thirteen types of antibiotic resistance genes (ARGs) and their associated genetic carriers (chromosome, plasmid, and unclassified) identified from the Nanopore datasets of presumptive CR bacteria isolated from nine wastewater samples. The numbers along the circumference of the diagram indicate the count of reads associated with each ARG and the carriers.

To gain a more detailed understanding of which specific subtypes of beta-lactamase genes were responsible for the observed resistance patterns in the presumptive CR bacteria, we analyzed the reads (454 reads in total) containing beta-lactamase genes (Fig. 5). A majority of these carbapenemase-associated genes, approximately 81.9%, were found to be on plasmids.

**Figure 5.**
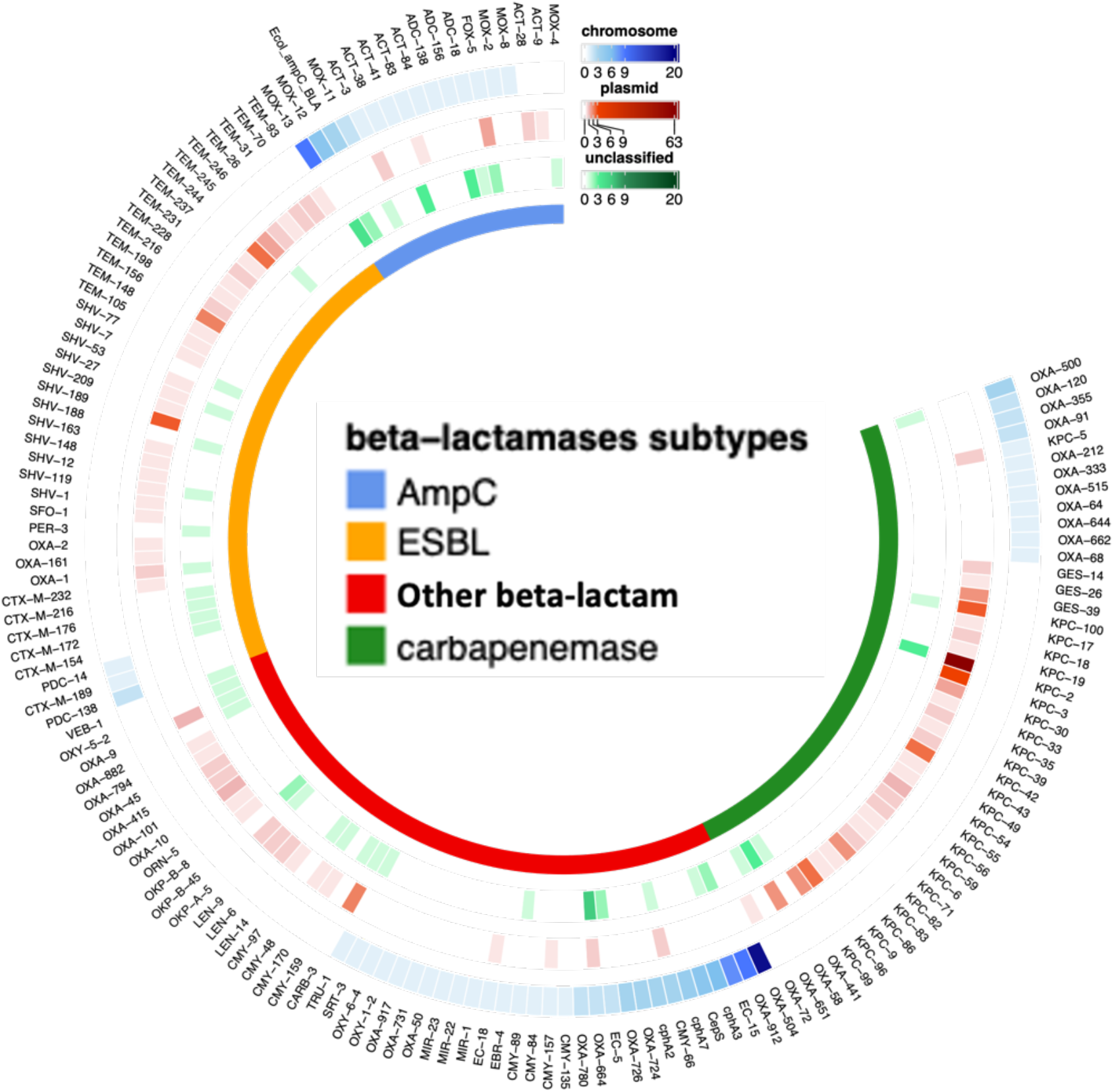
Distribution and abundance of antibiotic resistance gene (ARG) subtypes of beta-lactamase genes and frequency of associations with chromosomes, plasmids, or unclassified from the nine wastewater samples. The beta-lactamases are categorized into four groups: AmpC type, Extended-Spectrum Beta-Lactamases (ESBL), carbapenemase, and other beta-lactamase types. The heatmaps corresponding to the raw count of reads of chromosome-associated ARG subtypes are shown in blue, plasmid-associated ARG subtypes are shown in red, and unclassified ARG subtypes are shown in green.

Among the beta-lactamase genes, carbapenemase-encoding genes were the most prominent, accounting for 176 reads. This aligns with the primary mechanism of carbapenem resistance, which involves the production of carbapenemases. This category encompassed both Class A beta-lactamases (such as *bla*_KPC_) and Class D beta-lactamases, specifically, *bla*_OXA_. Consistent with the results from ddPCR, the *bla*_KPC_ family demonstrated the highest prevalence among the carbapenemase-encoding genes. Notably, within the *bla*_KPC_ family, the subtype *bla*_KPC-2_ was found to be the predominant gene. Our metagenomic sequencing results also revealed that some less commonly targeted carbapenemease-encoding genes (e.g., *bla*_GES_ family, *bla*_OXA-51-like_, *bla*_OXA-58-like,_ *bla*_OXA-211-like_), were found in the presumptive CR colonies, collectively accounting for 18.18% of all the detected carbapenemase-encoding reads.

In addition to carbapenemase-production, certain bacteria can develop carbapenem resistance due to a combination of factors, including reduced outer membrane permeability and the overproduction of β-lactamases, such as some AmpC-type or extended-spectrum β-lactamases (ESBL) that have limited carbapenem-hydrolyzing capabilities ^55^. We detected both ESBL and AmpC-type beta-lactamases in our samples, with a total of 130 reads associated with these two types of beta-lactamases. Among them, we observed a greater diversity of subtypes of ESBL genes (7 subtypes) as compared to AmpC-type genes (5 subtypes). Additionally, around 30% of the beta-lactamase reads corresponded to other beta-lactamase types, which are primarily associated with Class C beta-lactamases and confer resistance against different antibiotics, for example, cephalosporins and penams. While these beta-lactamases may have been co-selected by carbapenem, they were not classified as carbapenemases. Additionally, we identified many genes associated with multidrug efflux pumps. Previous studies have reported that, many Class C beta-lactamases exhibit a low activity for carbapenem hydrolysis when overproduced, and these enzymes can play a role in carbapenem resistance, particularly in conjunction with decreased outer-membrane permeability or increased efflux pump expression ^56^. This observation provides insight into the disparities observed between culture-based and ddPCR-based results as other carbapenem-resistance mechanisms beyond the five main carbapenemases commonly found in *Enterobacterales* may also play a role in the resistance phenotypes observed in cultured isolates. Given that the ddPCR assay solely targeted the five carbapenemase-encoding genes, drawing a direct comparison between these phenotypes and the five genes is inherently complex and likely overlooks many carbapenem resistance mechanisms present in the microbial community.

To further link the ARGs with their associated host, we used Centrifuge and found that only 25.6 % of the chromosomal reads carrying ARGs (266 reads) could be classified with taxonomic information (Fig. 6). The primary CR bacteria detected were species of *Pseudomonas, Enterobacterales*, and *Acinetobacter* (Fig. S6). These organisms, when exhibiting carbapenem resistance, are recognized as pathogens of substantial concern for public health, as emphasized in the 2019 AR Threats Report ^12^. In healthcare settings, the most frequently encountered carbapenemase-producing *Enterobacteriaceae* species among patients include *Escherichia coli* and *Klebsiella pneumoniae* ^57,58^. In sequencing the CR isolates, we found *Klebsiella pneumoniae*, and also environmental bacteria such as *Streptomyces* sp. with CR. Notably, most of the ARGs selected by carbapenem-supplemented culture-based method were associated with plasmids, and long-read sequencing alone cannot be used to confidently infer host information of plasmid-associated ARGs. Out of all the hosts identified with chromosomal ARGs, 39 of these ARGs were categorized as beta-lactamases. Within this group, four of the genes encoded carbapenemases, including *bla*_OXA-515_ (a subtype of *bla*_OXA-48-like_), *bla*_OXA-500_ (a subtype of *bla*_OXA-213-like_), and *bla*_OXA-355_ (a subtype of *bla*_OXA-229-like_). The remaining beta-lactamase genes consisted of 6 reads related to AmpC-type beta-lactamase and 13 reads associated with other OXA-type beta-lactamase.

**Figure 6.**
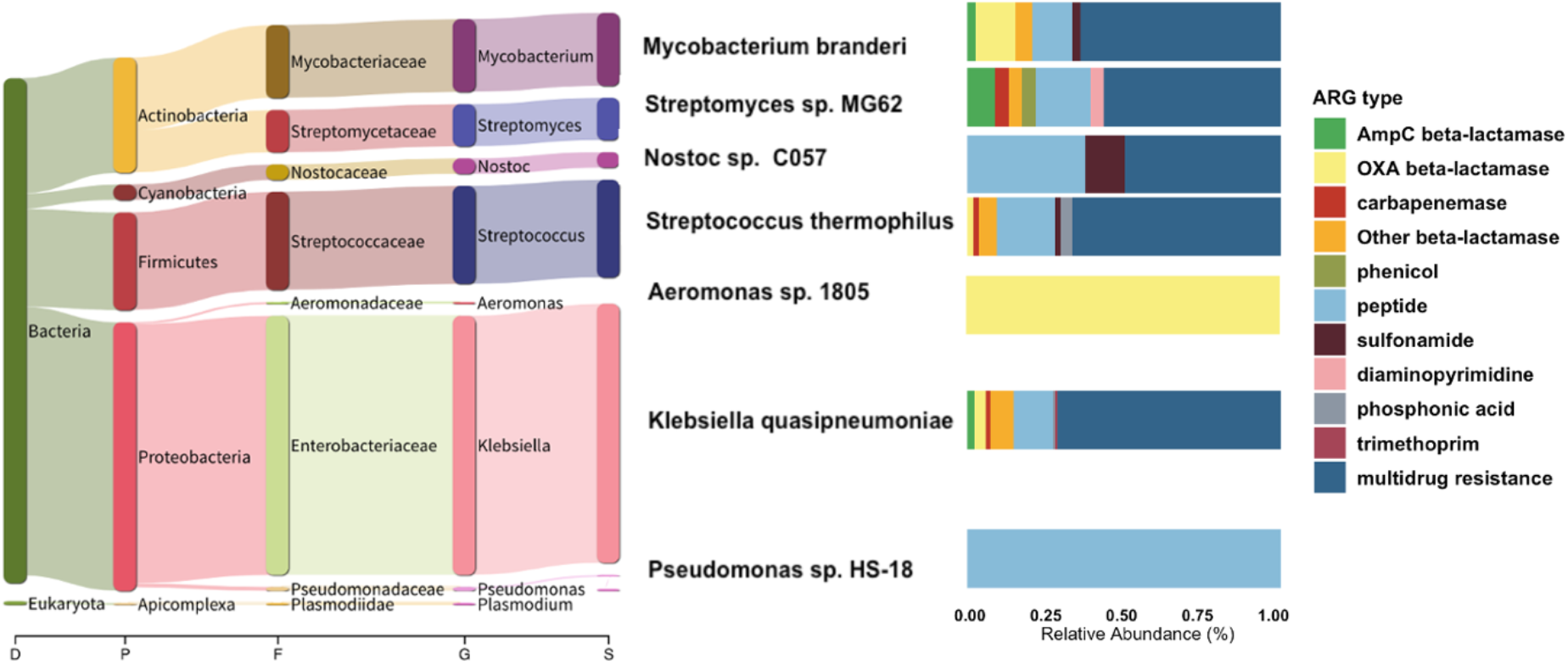
Bacterial host range of antibiotic resistance genes (ARGs) detected across all WWTP samples. The phylogenetic tree displays the composition and relative abundance of bacterial hosts of ARGs (the most abundant 7 species out of 15 are shown). The left panel bar chart illustrates the diversity and prevalence of ARGs carried by each corresponding species.

### 3.4 Limitations and Research Needs

One limitation of our study was its short duration, spanning only twelve weeks with weekly sampling. Consequently, the weak associations observed may be partly due to the limited time frame of our surveillance, which results in fewer data points for analysis. In the future, long-term surveillance may be needed to capture significant trends in population-level CR.

Furthermore, by sequencing the CR colonies, we were able to identify some clinically important species with CR resistance. However, only 15 species were identified from this chromosome-associated ARG dataset. A majority of the CR genes detected via sequencing were associated with plasmids. It is currently challenging to determine the bacterial hosts of plasmids via metagenomic sequencing due to their ability to cross species boundaries. In future studies, methods such as whole genome sequencing of isolates, single-cell sequencing, and proximity ligation sequencing methods (e.g., Hi-C) could be applied to identify plasmid-host relationships.

Despite this constraint, integrating sequencing with culture-based methods allowed us to validate the observed differences between ddPCR and culture-based methods. Therefore, careful consideration is necessary when selecting methods and comparing quantitative levels determined using different approaches. Moreover, wastewater monitoring should combine culture-based methods with sequencing to identify clinically important ARGs and their associated bacterial hosts. This combined approach can then be compared with clinical surveillance metrics, such as hospitalization rates and infections, to enhance the actionability of wastewater AR monitoring information.

## 4. Conclusion

In this study, we quantified and characterized carbapenem resistance in wastewater influent samples collected from three WWTPs. We evaluated and compared two commonly used methods for quantifying resistance: 1) selective culturing to measure carbapenem-resistant bacteria, and 2) ddPCR quantification of five carbapenemase-encoding genes. We demonstrated that both culture-based and ddPCR methods can provide quantitative levels of carbapenem resistance levels in wastewater. It is crucial, however, to cautiously interpret trends in AR levels when comparing data generated using different methods. Long-term, longitudinal monitoring of carbapenem resistance in wastewater is essential to produce actionable information. Combining culture-based methods with metagenomic sequencing (e.g., long-read sequencing, whole genome sequencing) of wastewater and clinical isolates can identify clinically important antibiotic resistance associated with hospital-acquired infections. This information is crucial for developing highly specific ddPCR assays that provide actionable insights. Moving forward, establishing high-throughput screening methods to elucidate the interactions between resistance genes and their bacterial hosts is critical for advancing AR wastewater monitoring and understanding of how resistance develops and spreads.

## Data Availability Statement

Sequencing data was deposited to the NCBI Sequence Read Archive under BioProject accession number: PRJNA1063622.

## Acknowledgements

This work was supported by the Centers for Disease Control and Prevention (Contract No. 75D30122C14709), the Centers for Disease Control and Prevention (ELC-ED grant no. 6NU50CK000557-01-05 and ELC-CORE grant no. NU50CK000557), and the Houston Health Department. We thank the efforts of the Rice wastewater monitoring team at Stadler lab for sample processing, including wastewater concentration and extraction. We also extend our gratitude to Houston Public Works for their assistance with sample collection and to the Houston Wastewater Epidemiology team for their contributions.

## Competing Interests

The authors declare that they have no known competing financial interests or personal relationships that could have appeared to influence the work reported in this paper.

## Notes

### Competing Interest Statement

The authors have declared no competing interest.

