## Supplementary materials for "Comparative analysis of culture- and ddPCR-based wastewater surveillance for carbapenem-resistant bacteria"

**1. Materials and Methods**

The protocols for wastewater sample processing, concentration, DNA extraction, and ddPCR quantification of gene targets are described as below.

**Influent solids concentration:** We aliquoted influent wastewater samples into two 50 mL conical centrifuge tubes. We thoroughly homogenized wastewater samples by shaking sample bottles prior to and in between aliquoting each replicate. 50 mL wastewater samples were centrifuged for 20 minutes at 4,100 RPM and 4 °C. After centrifugation, we carefully poured off the supernatant and saved the solid portion of sample leftover in the centrifuge tubes and added 1 mL lysis buffer to each centrifuge tube and mixed well with pipette in order to suspend the solids pellet. Once suspended, all content from the centrifuge tube was transferred into a 2 mL bead beating tube.

**Extraction:** DNA extraction was performed using a Chemagic™ Prime Viral DNA/RNA 300 Kit H96 (Chemagic, CMG-1433, PerkinElmer). We followed the manufacturer's recommended protocol for the sample preparation. The tubes were bead beaten at max speed in a Mini-Beadbeater 24 (3,500 RPM; 112011, BioSpec) for 1 minute, set on ice for 2 minutes, and bead beaten again at max speed for 1 minute. After bead beating, the tubes were centrifuged to pellet the beads (17,000 g, 4 °C). Subsequently, 300 µL of supernatant from each bead tube was loaded into a 96-deep well plate followed by addition of 300 µL lysis buffer and 14 µL of Proteinase K as directed by the manufacturer's protocol. Apart from the sample plate, an elution buffer plate, a magnetic bead plate, a wash buffer plate, and an eluate collection plate were prepared as directed by the protocol. All plates were loaded onto the Chemagic. The extraction program "Chemagic

Viral300 360 H96 drying prefilling VD200309.che" was selected for automated nucleic acid extraction. Each sample was eluted to generate a 50 µL nucleic acid extract. After nucleic acid extraction, all sample extracts were sealed and stored at - 20 °C for ddPCR analysis.

**Commented [LS1]:** Not true?

**Quantification:** ddPCR was performed on a QX600 AutoDG Droplet Digital PCR System (Bio-Rad) and a C1000 Thermal Cycler (Bio-Rad) in 96-well optical plates. Five carbapenemase-encoding gene (*bla*$_{IMP}$, *bla*$_{NDM}$, *bla*$_{VIM}$, *bla*$_{OXA\text{-}48\text{-}like}$ and *bla*$_{KPC}$) targets were quantified in wastewater samples using a multiplex ddPCR assay. Briefly, a 22 µl reaction mix containing 10 µl of nucleic acid was mixed with the ddPCR Multiplex Supermix (Bio-Rad) following the manufacturer's protocol. Reaction mix compositions and thermal cycling conditions are detailed in Tables (S2-S5). Droplets were read on a QX600 Droplet Reader (Bio-Rad) and analyzed using QuantaSoft v1.7.4 software. Droplets were manually thresholded per channel and data were exported to an Excel file for further analysis.

**LOD:** A limit of detection (LOD) ≥ 3 positive droplets and an acceptable total generated droplet count of at least 10,000 were established for all sample wells as recommended by the manufacturer, and based on methods previously described by Lou et al., 2022 [1]. In addition to the 3 droplets threshold, the initial LOD concentration for the plate was calculated by averaging the concentrations of all negative control samples on a given plate. A copy number of 0.7 gene copies/µL was assigned to any plate when none of the negative controls contained any positive droplets. This corresponds to the concentration of 3 positive droplets among 10,000 total generated droplets.

**Table S1. Wastewater treatment plants sampled, service populations, and geographic service areas.**

| Wastewater treatment plant | Abbreviation | Population | Average gal/cap/day | Area, square miles |
|---|---|---|---|---|
| 69th Street | 69 | 551,150 | 145 | 96.72 |
| Upper Brays | UB | 97,918 | 105 | 12.81 |
| West District | WD | 85,129 | 118 | 17.86 |

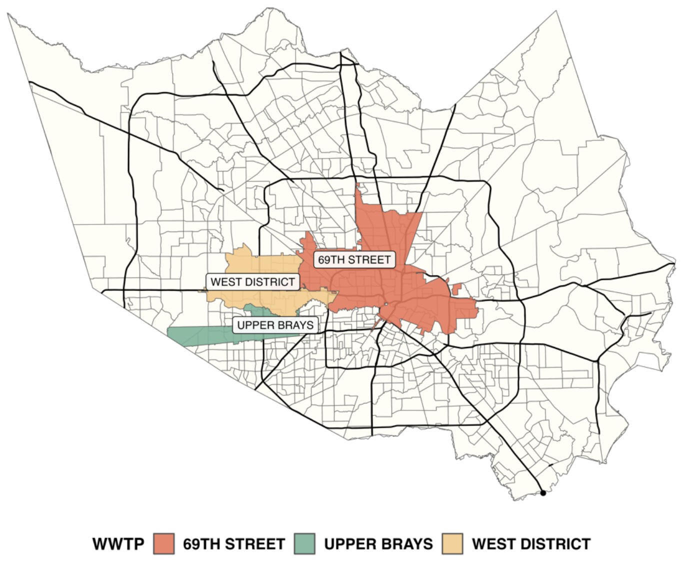


**Figure S1. Service areas of the three wastewater treatment plants (69, UB, WD).**

**Table S2. Concentration factors for sample processing procedure.**

| Concentration and extraction procedures | Vol. | Unit | Concentration Factor |
|---|---|---|---|
| Raw wastewater samples for centrifugation | 50 | ml | - |
| Sample with lysis buffer for bead-beating | 1000 | ul | 50 |

| | | | |
|---|---|---|---|
| Lysate transferred after bead-beating | 300 | ul | |
| Mastermix added [lysis buffer, Poly(A) RNA and proteinase K] for Chemagic | 314 | ul | |
| DNA Elution volume | 50 | ul | 6 |
| Overall concentration factor | | | 300 |

**Table S4. Primer and probe sequences used for ddPCR quantification**

| **Targets** | **Primer/probe** | **Sequence (5'-3')** | **Amplicon size (bp)** |
|---|---|---|---|
| ***bla***$_{\textbf{OXA-48-like}}$ | Oxa-f | TTACCC GCATCT ACCTTT | 186 |
| | Oxa-r | GCGGG CAAATT CTTGAT A | |
| | Oxa-p | /5Cy55/TG GTT AAG GAT GAA CAC CAA GTC TT/3IAbRQSp/ | |
| ***bla***$_{\textbf{VIM}}$ | Vim-f | YAATG GWCTC ATTGTC CGTG | 80 |
| | Vim-r | GAAGT GCCGC TGTGTT TTTC | |
| | Vim-p | /5Cy5/CT TYT KAT T/TAO/G ATA CAG CGT | |

| | | | |
|---|---|---|---|
| | | GGG<br>GTG<br>C/3IAbR<br>QSp/ | |
| ***bla*$_{KPC}$** | Kpc-f | CGGAA<br>CCATTC<br>GCTAA<br>ACTC | 135 |
| | Kpc-r | GAAAG<br>CCCTTG<br>AATGA<br>GCTG | |
| | Kpc-p | /56-<br>ROXN/A<br>C TTT<br>GGC<br>GGC<br>TCC<br>ATC<br>GGT<br>GTG<br>TA/3IAb<br>RQSp/ | |
| ***bla*$_{IMP}$** | Imp-f | AGTYA<br>MTTGG<br>TTTGTG<br>GAGC | 93 |
| | Imp-r | TTAAG<br>CCACT<br>CTATTC<br>CNCC | |
| | Imp-p | /56-<br>FAM/CC<br>TCW<br>CAT<br>T/ZEN/T<br>YCA<br>TAG<br>CGA<br>CAG<br>CAC<br>G/3IABk<br>FQ/ | |
| ***bla*$_{NDM}$** | Ndm-f | GATTGC<br>GACTT | 189 |

| | | |
|---|---|---|
| | | ATGCC AATG |
| | Ndm-r | TCGATC CCAAC GGTGA TATT |
| | Ndm-p | /5SUN/A C ACA GCC T/ZEN/G ACT TTC GCC G/3IABk FQ/ |

**a.**

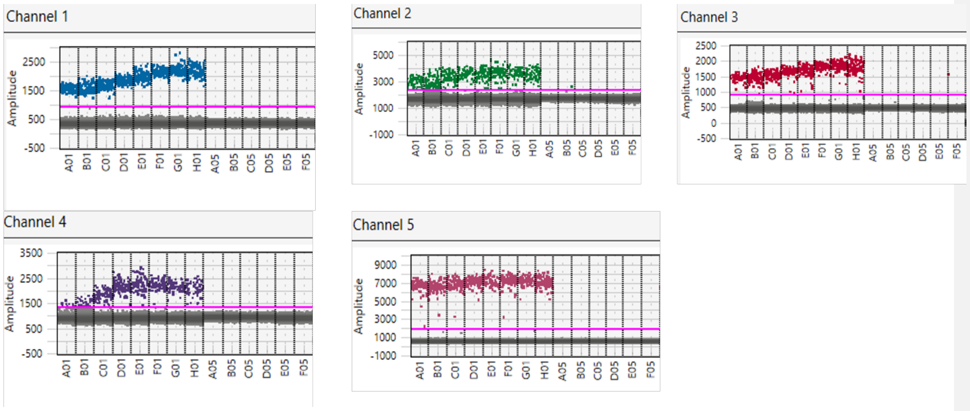


**b.**

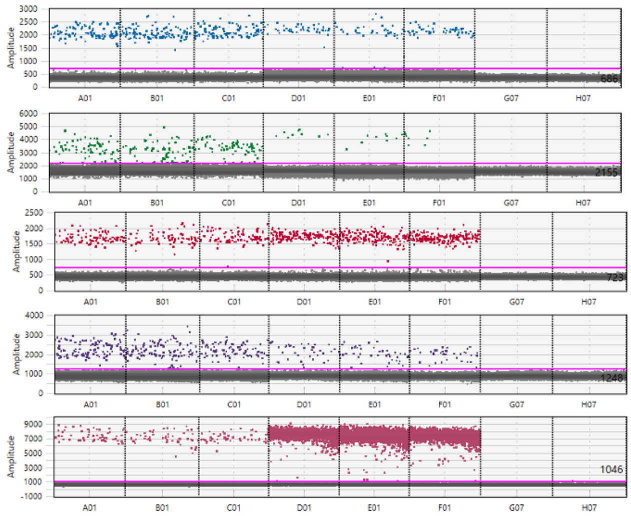


**Figure S2.** Multiplexed ddPCR assay optimization and application. (a) Annealing temperature optimization across a gradient from 56.2 °C to 62 °C (wells A to H) to determine optimal conditions. Gblocks were distributed across wells A01 to H01, with no-template controls (NTCs) positioned in wells A05 to F05. (b) Implementation of the multiplexed ddPCR assay using gblocks (wells A01 to C01), wastewater samples (wells D01 to F01), and additional NTCs (wells G07 to H07) to assess the presence of five genes. The targets, visualized in channels 1 to 5, include *bla*$_{\text{IMP}}$ (FAM, depicted as blue dots), *bla*$_{\text{NDM}}$ (SUN, green dots), *bla*$_{\text{VIM}}$ (Cy5, red dots), *bla*$_{\text{OXA-48-like}}$ (Cy5.5, purple dots), and *bla*$_{\text{KPC}}$ (ROX, pink dots).

**Table S5. Reaction composition for multiplex ddPCR assay.**

| Component | Final concentration |
|---|---|
| Multiplex Supermix | 1x |
| Primer/Probe - *bla*$_{\text{IMP}}$ | 900 nM/ 250 nM |
| Primer/Probe - *bla*$_{\text{NDM}}$ | 900 nM/ 250 nM |
| Primer/Probe - *bla*$_{\text{VIM}}$ | 900 nM/ 250 nM |
| Primer/Probe - *bla*$_{\text{OXA-48-like}}$ | 900 nM/ 250 nM |

| | |
|---|---|
| Primer/Probe - *bla*KPC | 900 nM/ 250 nM |
| RNase/DNase free water | variable |
| DNA template | variable |

**Table S6. Thermal cycling conditions for multiplex ddPCR assay.**

| Cycling Step | Temperature °C | Time | Number of Cycles |
|---|---|---|---|
| Enzyme activation | 95 | 10 min | 1 |
| Denaturation | 95 | 1min | 40 |
| Annealing/Extension | 57.9 | 1min* | |
| Enzyme Deactivation | 72 | 2 min | |
| Droplet Stabilization | 4 | 30 min | 1 |
| Hold | 4 | ∞ | |

*Ramp rate is set to 2 °C / sec.

**Table S7. Five gene fragment standards (gblocks) as positive controls.**

| gene | gblock |
|---|---|
| *bla*KPC | CGGAACCATTCGCTAAACTCGAACAGGACTTTGGCGGCTCCAT CGGTGTGTACGCGATGGATACCGGCTCAGGCGCAACTGTAAGT TACCGCGCTGAGGAGCGCTTCCCACTGTGCAGCTCATTCAAGG GCTTTC |
| *bla*IMP | TTCCTAAACATGGTTTGGTGGTTCTTGTAAATGCTGAGGCTTAC CTAATTGACACTCCATTTACGGCTAAAGATACTGAAAAGTTAG TCACTTGGTTTGTGGAGCGTGGCTATAAAATAAAAGGCAGCAT TTCCTCTCATTTTCATAGCGACAGCACGGGCGGAATAGAGTGG CTTAATTCTCGATCTATCCCCACGTATGCATCTGAATTAACAAA TGAACTGCTTAAAAAAGACGGTAAGGTTCAAGCCACAAATTC ATTTAGCGGAGTTAACTATTGGCTAGTTAAAAATAAAATTGA |
| *bla*VIM & *bla*NDM | GTTTGGTCGCATATCGCAACGCAGTCGTTTGATGGCGCGGTCT ACCCGTCCAATGGTCTCATTGTCCGTGATGGTGATGAGTTGCTT TTGATTGATACAGCGTGGGGTGCGAAAAACACAGCGGCACTTC TCGCGGAGATTGAAAAGCAAAGATTGCGACTTATGCCAATGC GTTGTCGAACCAGCTTGCCCCGCAAGAGGGGATGGTTGCGGCG CAACACAGCCTGACTTTCGCCGCCAATGGCTGGGTCGAACCAG CAACCGCGCCCAACTTTGGCCCGCTCAAGGTATTTTACCCCGG CCCCGGCCACACCAGTGACAATATCACCGTTGGGATCGACGGC ACCG |
| *bla*OXA-48-like | ATTTTTACCCGCATCTACCTTTAAAATTCCCAATAGCTTGATCG CCCTCGATTTGGGCGTGGTTAAGGATGAACACCAAGTCTTTAA GTGGGATGGACAGACGCGCGATATCGCCACTTGGAATCGCGA |

| | |
|---|---|
| | TCATAATCTAATCACCGCGATGAAATATTCGGTTGTGCCTGTTT ATCAAGAATTTGCCCGCCAAATTGGCGAGGCACGTATGAGCA AGATGCTACATGCTTTCGATTATGGTAATGAGGACATTTCGGG CAATGTAGATACTTTTTGGCTTGATGGTGGTATTCGAATTTC |

### 2. Results and Discussion

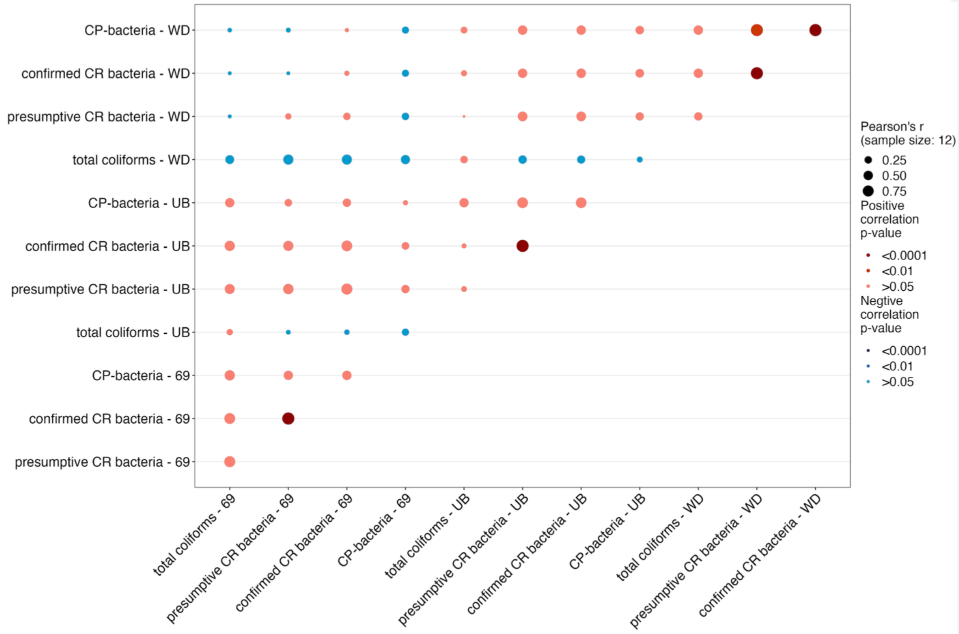


**Figure S3.** Pearson correlation of concentrations of total coliforms, presumptive CR bacteria, confirmed CR bacteria, and carbapenemase-producing bacteria within and across WWTPs. Bubble size reflects the Pearson correlation coefficient, with larger bubbles representing stronger correlations. The color of the bubble indicates the p-value's significance level: blue signifies a negative correlation, red signifies a positive correlation, and darker shades indicate stronger p-values.

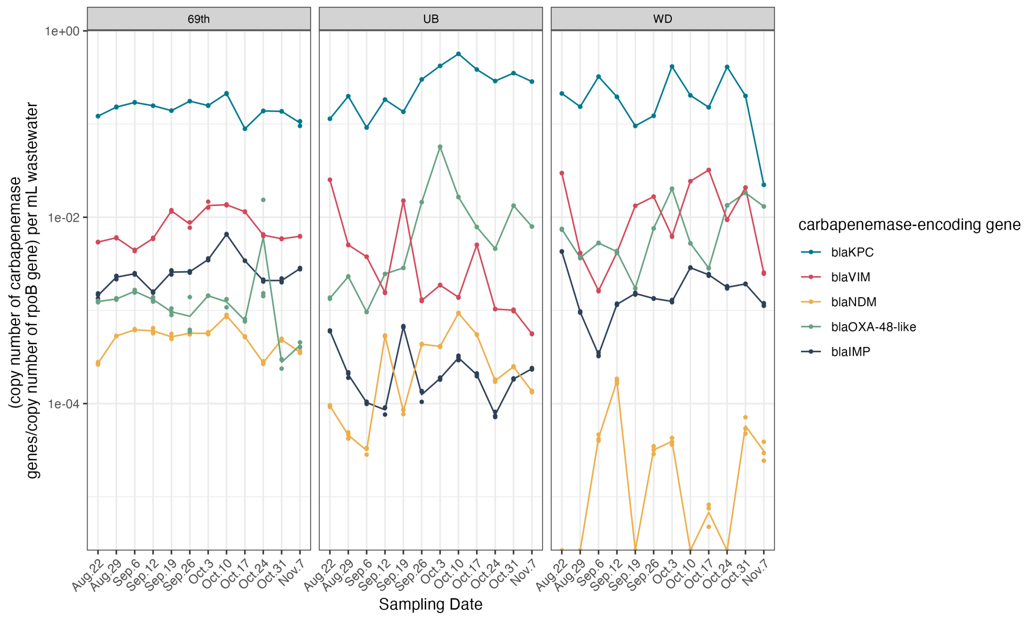


**Figure S4.** Relative abundance of carbapenemase-encoding genes (copy number of ARG/copy number of *rpoB* genes per mL) in influent over 12 weeks from three WWTPs. Each panel represents a different WWTP. Each dot represents a replicate measurement, and the lines connect the mean values of the replicates.

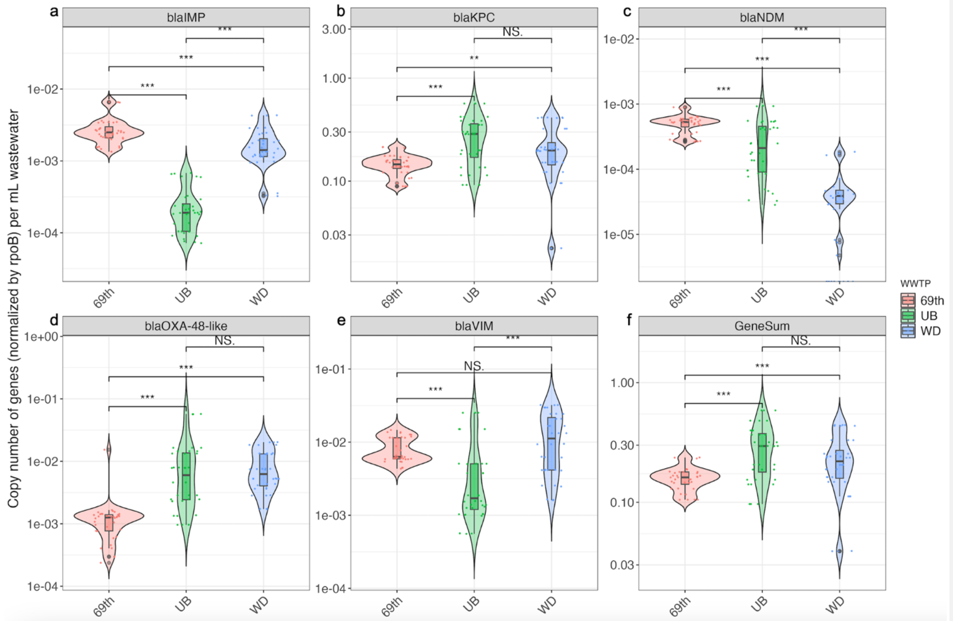


**Figure S5.** Comparison of carbapenemase gene relative abundances (normalizaed by *rpoB* gene) across WWTPs (panels a – e) and cumulative concentration of the five genes (panel f). Wilcoxon rank-sum test was used to determine statistical significance. Violin plots depict the distribution of gene copy numbers within each WWTP, with dots symbolizing individual data points. Inner boxes delineate the dataset's quartiles, with solid lines marking the medians. The whiskers show the minimum and maximum values. The asterisks represent the significance levels, '*' for $p < 0.05$, '**' for $p < 0.01$, '***' for $p < 0.001$, and 'NS' means not significant or $p > 0.05$.

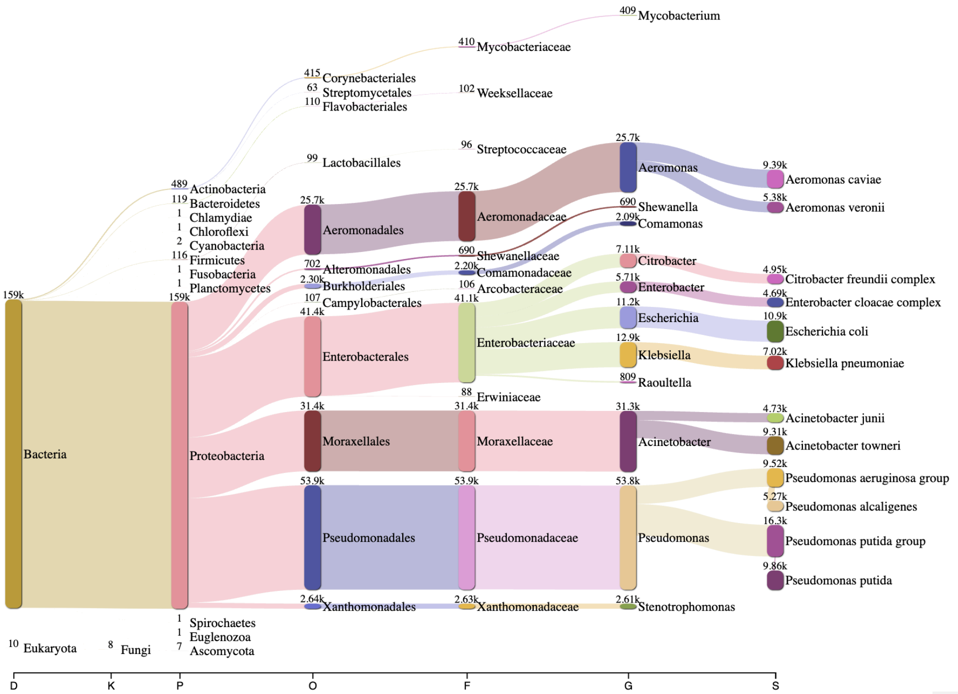


**Figure S6.** Distributions of presumptive carbapenem-resistant bacteria from all influent samples collected from three WWTPs. The phylogenetic tree displays the diversity and relative abundance of presumptive carbapenem-resistant bacteria isolated using culture-based methods. The numbers displayed indicate the read counts. All chromosomal reads over 1,000 bp underwent taxonomic classification.
